# Interstitial macrophages prevent tuberculosis relapse by restricting *Mycobacterium tuberculosis* immune evasion

**DOI:** 10.1101/2025.08.27.672444

**Authors:** V Vinette, A Castro, H Kim, C Trujillo, M Xie, M Gengenbacher, TR Ioerger, S Ehrt

## Abstract

Alveolar macrophages (AMs) are the first immune cells to encounter *Mycobacterium tuberculosis* (Mtb) in the lungs, but they frequently fail to eliminate this causative agent of tuberculosis (TB), allowing Mtb to persist or replicate. Interstitial macrophages (IMs) are recruited to restrict Mtb growth and limit immune evasion. While IMs have been implicated in the control of acute Mtb infection, their role during latent tuberculosis infection (LTBI) has not yet been explored. We hypothesized that IMs contribute to maintaining latency and that their depletion during LTBI would promote Mtb reactivation, leading to TB relapse and disease. To test this, we utilized our previously established mouse model of paucibacillary Mtb infection that mimics aspects of LTBI in humans to selectively deplete IMs during the latent phase. IM depletion led to TB relapse in 26% of mice compared to 2% in control mice. The transitory depletion of this macrophage subset transiently affected both pulmonary macrophage and neutrophil populations. Mice that relapsed exhibited an increased proportion of pro-inflammatory IMs and elevated concentrations of G-CSF, GM-CSF, IL3, IL-12, IL-13, IL-17A and KC in the lung. These findings indicate that IMs play a critical role in controlling latent Mtb and preventing TB relapse.

## 1. INTRODUCTION

Tuberculosis (TB) is the deadliest infectious disease in the world, affecting 10.8 million people annually and causing 1.3 million deaths in 2023 (WHO, 2024). Approximately 95% of people exposed to Mtb do not develop the disease. Some individuals develop latent TB infection (LTBI), where the bacteria persist in the body without causing any clinical symptoms. Sometimes, the pathogen reactivates and causes active TB. *Mycobacterium tuberculosis* (Mtb), the causative agent of this disease, is first encountered in the lung by tissue resident alveolar macrophages (AMs). While AMs are the initial line of defense, they are not always able to eliminate intracellular Mtb, allowing the bacteria to persist and potentially disseminate. Interstitial macrophages (IMs), on the other hand, are located within the lung interstitium and are recruited to the site of infection to control Mtb and restrict their evasion from the immune system. IMs are more pro-inflammatory in nature (Pisu et al., 2020, 2021) and more restrictive for Mtb than AMs (Huang et al., 2018). In a mouse model of acute TB, IM depletion during the early stage of infection promoted the growth of Mtb in the lung, highlighting their importance in the early immune response to Mtb infection (Huang et al., 2018). The cellular immune control of LTBI is multifactorial and depends on a delicate balance from the host immune system to prevent Mtb reactivation without causing excessive inflammation. Notably, the formation and maintenance of the granuloma throughout the infection have been shown to be crucial in this process (Dutta et al., 2014; Flynn et al., 2011a; McCaffrey et al., 2022; Silva Miranda et al., 2012). It is composed mainly of macrophages as the core but also contains T cells and other immune cells (Flynn et al., 2011a; McCaffrey et al., 2022). Macrophages provide the cellular niche for Mtb, where the pathogen can persist in a non-replicating or slowly replicating state (Flynn et al., 2011b; McCaffrey et al., 2022). T cell-mediated control from both CD4^+^ and CD8^+^ cells is also important in controlling Mtb, in part by activating infected macrophages (Adekambi et al., 2015; Hou et al., 2021). However, the role of interstitial macrophages specifically in controlling latent Mtb has yet to be explored.

To assess the importance of IMs for Mtb control during latency, we used a paucibacillary mouse model that mimics aspects of human LTBI and yields reproducible relapse frequencies across independent experiments (H. Su et al., 2021). We have previously shown that transient genetic depletion of the essential Mtb protein biotin protein ligase (BPL) leads to apparent sterilization of Mtb and induces a state of paucibacillary Mtb infection in mice, followed by spontaneous relapse in about 30% of mice ten months post-infection (H. Su et al., 2021). Considering that macrophages are the main reservoir for Mtb and that IMs are important in restricting Mtb during acute infection, we hypothesized that depleting IMs during the paucibacillary, latent phase of Mtb infection would result in a resurgence of Mtb growth and consequently increased TB relapse. Here, we demonstrate that transient IM depletion during latency promotes TB relapse in mice. Longitudinal cytokine expression analyses and flow cytometry immunoprofiling performed in parallel revealed an increased frequency of pro-inflammatory IMs and an altered cytokine milieu in mice with TB relapse. These data indicate that IMs are critical for restraining persistent Mtb, maintaining them in latency, and consequently limiting TB reactivation and disease.

## 2. MATERIALS AND METHODS

### Mtb strains and culturing conditions

Mtb H37Rv BPL-DUC has been previously described (Su et al., 2021; Tiwari et al., 2018). Strains were cultured in liquid Middlebrook 7H9 medium supplemented with 0.2% glycerol, 0.05% tyloxapol and ADNaCl (0.5% BSA, 0.2% dextrose and 0.085% NaCl), and on Middlebrook 7H10 agar plates supplemented with 0.2% glycerol and Middlebrook OADC enrichment. Antibiotics were added to select for the genetically modified strain as described (Su et al., 2021).

### Mouse infections

8-10-week old female C57BL/6 mice were infected with Mtb BPL-DUC using an inhalation exposure system (Glas-Col) with Mtb in a mid-log growth phase to deliver ∼100 bacilli per mouse. Mice received doxycycline-containing chow (2,000 ppm, Research Diets, Inc.) for the indicated period. Animals were housed in a biosafety level 3 (BSL3) facility and fed water and chow *ad libitum*. All procedures involving animals were reviewed approved by the Weill Cornell Medicine Institutional Animal Care and Use Committee (IACUC).

### Enumeration of bacterial burden

To enumerate CFUs, organs were homogenized in PBS and cultured on 7H10 agar. To recover Mtb BPL-DUC from mice that were treated with doxycycline, charcoal (0.4% wt/vol) was added to the plates. Agar plates were incubated for at least four weeks at 37°C.

### Depletion of interstitial macrophages

Control liposomes (PBS-loaded) and clodronate-loaded liposomes were purchased from LIPOSOMA research. To deplete interstitial macrophages, mice were injected intravenously with 200 μL clodronate liposomes during the paucibacillary phase every 3-4 days over a period of two weeks for a total of four injections. PBS-loaded liposomes were used as controls. Efficiency of macrophage depletion was assessed by flow cytometry.

### Flow cytometry

For flow cytometry, mouse lungs and spleens were isolated as previously described (H. Su et al., 2021), with some modifications. Briefly, lungs were isolated in digestion media containing 0.5% BSA, and 150 U/mL collagenase IV and 100 μg/mL DNase I enzymes in C-tubes (Miltenyi Biotec). Lungs were homogenized using the GentleMACS Tissue Dissociator using the program *m_lung_02.01* and then incubated at 37°C for 45 minutes. Cells were homogenized one more time using the program *m_lung_01_02* and filtered using 70 μm cell strainers, collected by centrifugation at 500x*g* for 5 minutes and resuspended in 3 mL red blood cell lysis buffer (eBiosciences). Cells were incubated for 5 minutes at room temperature with occasional shaking. PBS was added to the tube, centrifuged and resuspended in RPMI 1640 media. Spleens were harvested in PBS in C-tubes and homogenized using the GentleMACS Tissue Dissociator. Cells were filtered using 40 μm cell strainers, collected by centrifugation at 500x*g* for 5 minutes and resuspended in 3 mL red blood cell lysis buffer and processed similarly to lung cells. Samples were seeded in a 96-well plate and kept at 4°C overnight. Cells were then stained with Zombie UV LIVE/DEAD Fixable Blue stain (BioLegend) to discriminate live and dead cells. Purified anti-CD16/32 antibody (clone 93) was used to block Fc receptors on all cells before staining. The antibody cocktail was prepared in a 3:1 solution of Cell Staining Buffer and Brilliant Stain Buffer and cells were stained for 30 minutes at 4°C with the following anti-mouse antibodies: CD45-AF700 (clone 30-F11), CD11b-BUV395 (clone M1/70), CD11c-FITC (clone N418), Ly6C-BV785 (clone HK1.4), Ly6G-APC (clone 1A8), CD62L-BUV737 (clone MEL-14), CD63-PE-Cy7 (clone NVG-2), Siglec-F-BV421 (clone S17007L), CX3CR1-PE (clone SA011F11), I-A/I-E-BB700 (clone 2G9), CD38-PE/Dazzle594 (clone 90), CD103-BV711 (clone M290) and CD24-BV650 (clone M1/69). The cells were then fixed with fixation buffer (BioLegend) for 30 minutes at room temperature and washed twice with cell staining buffer before being and taken out of the BSL3. Samples were acquired using the BD FACSymphony A5 cytometer and analyzed with FlowJo v10.

### Reactive oxygen species assay

To measure the levels of reactive oxygen species (ROS) in mice, blood was collected from the tail vein of mice into an anti-coagulant EDTA tube. Cells were transferred to a round-bottom 96-well plate, centrifuged at 500x*g* for 5 minutes and resuspended in ROS media (RPMI plus 10% FBS, 1 mM MgCl_2_ and 1 mM CaCl_2_). Cells were stimulated *ex vivo* with 100 ng/mL PMA at 37°C for 30 minutes. 5 μM CellROX Green (Thermo Fisher Scientific) was added to each sample and cell were incubated at 37°C for 30 minutes, after which cells were centrifuged and washed three times in PBS. Fc block was diluted in FACS-ROS buffer (PBS plus 2% heat-inactivated FBS and 2 mM EDTA) and added to all samples and cells were incubated at 4°C for 10 minutes. After centrifugation and removing the supernatant, cells were incubated with an antibody cocktail prepared in FACS-ROS buffer with antibodies CD45-AF700, CD11b-BUV395 and Ly6G-APC to identify neutrophils. The rest of the flow cytometry staining was performed as aforementioned. Before acquiring the samples on a BD FACSymphony A5 cytometer, cells were resuspended in 200 μL FACS-ROS buffer.

### Multiplex cytokine analysis

The concentrations of various cytokines and chemokines in the lung homogenates were measured using the Bio-Plex Pro Mouse Cytokine Group I 23-plex Assay (Bio-Rad) according to manufacturer’s protocol and as described previously (Tsai et al., 2024). Briefly, 50 μL of coupled beads were added to each well of a black, 96-well flat-bottom plate and washed twice with 100 µL of Bio-Plex wash buffer. Samples were diluted 1:2 in Bio-Plex sample diluent containing 0.5% (w/v) bovine serum albumin. Next, 50 µL of diluted samples, standards, and blanks (diluent + BSA) were added to the wells and incubated at room temperature for 30 min with shaking (850 rpm) in the dark. Plates were washed three times and incubated with 25 µL of 1X detection antibodies for 30 minutes with shaking in the dark. After another three washes, 50 µL of 1X Streptavidin-PE was added and incubated at room temperature for 10 minutes with shaking in the dark. Following three final washes, beads were resuspended in 125 µL of Bio-Plex assay buffer and analyzed using FLEXMAP 3D system (Millipore) according to the manufacturer’s instructions. Raw data were processed and analyzed using Belysa software (version 1.2.1, Millipore).

### Statistical analyses

Flow cytometry data were analyzed by t-test, or by one-way or two-way ANOVA with Tukey’s multiple comparisons test, depending on the experiment.

The cytokine measurements for days 1, 28 and 180 (**Figure 1F**) and day 194 (**Figure 2K**) were log-transformed to accommodate the wide variation of scales across various cytokines. These were analyzed by two-way ANOVA with a Tukey’s multiple comparisons test. Cytokine measurements for endpoint at day 294 (**Figure 3D**) were analyzed by fitting a linear-mixed model with random effects and applying an F-test to evaluate the significance of coefficients in the model. 11 cytokines (Cyt) were included, and three treatment groups (Tx) were used: control liposomes, clodronate-treated/non-relapsed, and clodronate-treated/relapsed. The Tx*Cyt interaction term incorporated distinct means for each cytokine in each treatment group. Since 4 of the 22 mice had relapsed, resulting in higher and more variable measurements, the relapsed mice were indicated with a binary covariate (Rpd) and the coefficients were treated as random effects for each mouse, allowing for different magnitudes of increase for each mouse. The data for all cytokines, mice, and controls was fit using *lmer()* in R, using the following formula: *log(obs) ∼ Tx*Cyt + (Rpd|Mouse)*. After fitting the model, all 3 pairwise contrasts among treatments were compared for each cytokine using an F-test using *emmeans()* in R. P values were adjusted for multiple comparisons using Tukey’s method (Tukey, 1949). Cytokines exhibiting significant differences (p value < 0.05) between clodronate/relapsed and control, or clodronate/relapsed and clodronate/non-relapsed, were identified.

**Figure 1.**
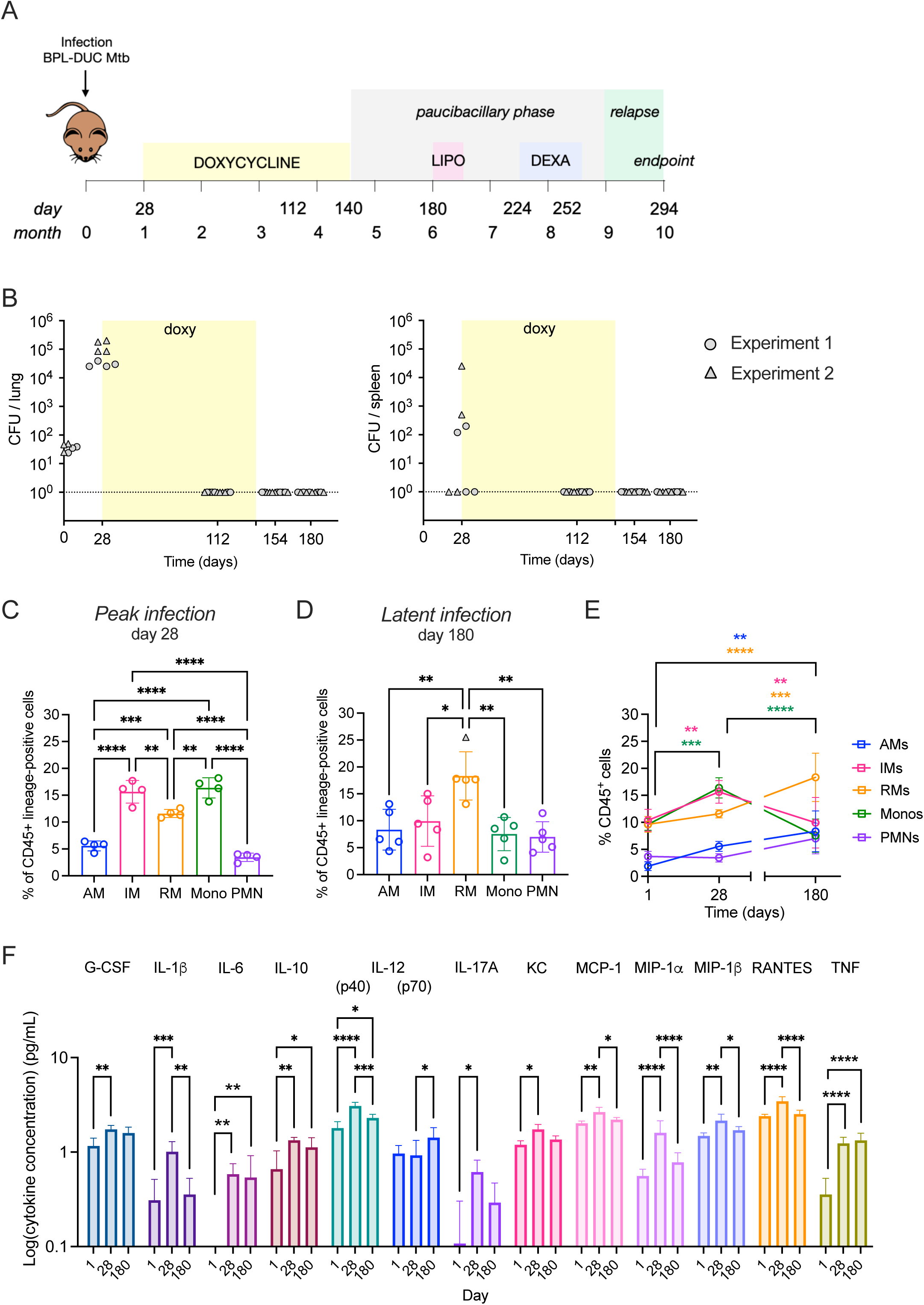
The pulmonary myeloid cell compartment and cytokine milieu are altered in a paucibacillary mouse model of latent tuberculosis infection. (A) Study design. Mice were infected with Mtb BPL-DUC by aerosol. Doxycycline (yellow) was administered for 16 weeks to deplete BPL, kill the bacteria and attain the paucibacillary phase (gray). Chlodronate liposomes or control liposomes (lipo) were injected intravenously every two or three days for two weeks to deplete interstitial macrophages (IMs) (pink). One month after treatment, dexamethasone (dexa) was administered to a group of mice treated with control liposomes for one month (blue). Relapse was assessed at three months post IM depletion on days 287-302, with 294 depicted as mean. (B) Bacterial burden was assessed throughout the infection by CFU determinations in the lung and spleen of mice. Different symbols (circle, triangle) denote data from two independent experiments that were combined for analysis. (C, D) Immunoprofiling was performed by flow cytometry with lung homogenates to assess the frequency of alveolar macrophages (AM), interstitial macrophages (IM) recruited monocytes/macrophages (RM), monocytes (Mono) and neutrophils (PMN) at the peak of infection (C) and during the paucibacillary phase, prior to treatment with liposomes (D). (E) Frequency of macrophage, monocyte and neutrophil populations in the lung over time before treatment with liposomes. (F) Cytokine concentrations in the lung of mice at days 1, 28 and 180. Data represent the mean ± SD of four or five biological replicates. * p < 0.05, ** p < 0.01, *** p < 0.001, **** p < 0.0001 based on one-way ANOVA (B and C) or two-way ANOVA (D and E) statistical test with Tukey’s multiple comparisons tests.

**Figure 2.**
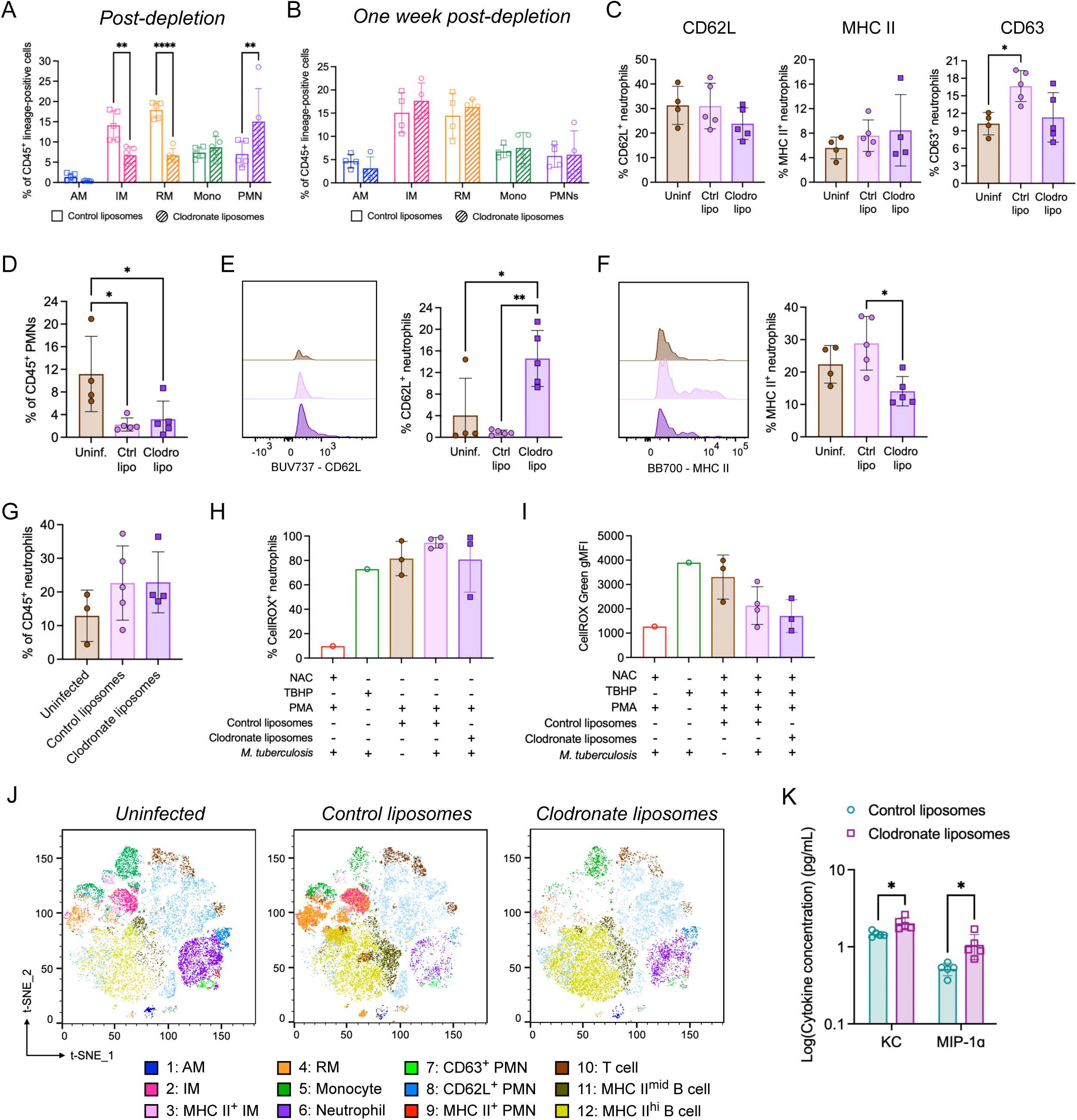
Clodronate liposome treatment during LTBI leads to a depletion of interstitial macrophages and increased recruitment of neutrophils to the lung. Mice were treated with control or clodronate liposomes for two weeks during the paucibacillary phase, and lung (A-C), spleen (D-F) and blood (G-I) were analyzed. (A-B) Immunoprofiling was performed by flow cytometry with lung homogenates to assess the frequency of alveolar macrophages (AM), interstitial macrophages (IM), recruited monocytes/ macrophages (RM), monocytes (Mono) and neutrophils (PMN) immediately post-depletion with liposomes (A) and one-week post-depletion (B). (C) Flow cytometry from lung homogenates following liposome treatment was done to assess the frequency of neutrophils expressing the activation markers CD62L and MHC II, and the degranulation marker CD63 in uninfected mice (Uninf) and mice treated with control liposomes (Ctrl lipo) or clodronate liposomes (Clodro lipo). (D-F) Flow cytometry from spleen homogenates following liposome treatment was performed to assess the frequency of neutrophils (D) and the fraction of these neutrophils expressing the activation markers CD62L (E) or MHC II (F). (G) Blood was collected from mice after IM depletion to assess the frequency of circulating neutrophils. (H-I) ROS assay performed with blood to assess the frequency of ROS-producing neutrophils (H) and the geometric mean fluorescence intensity (gMFI) of ROS production from neutrophils (I). (J) t-SNE plots of cell subtypes after clustering, colored according to cellular identity and separated by condition immediately following treatment, namely uninfected mice, and infected mice treated with control or clodronate liposomes. (K) Quantification of cytokine concentrations in the lung of mice treated with control or clodronate liposomes immediately following liposome treatment. Data represents the mean ± SD of three to five biological replicates. * p < 0.05, ** p < 0.01, **** p < 0.0001 based on one-way ANOVA (for C-I) or two-way ANOVA (for A, B and K) statistical test with Tukey’s multiple comparisons test.

**Figure 3.**
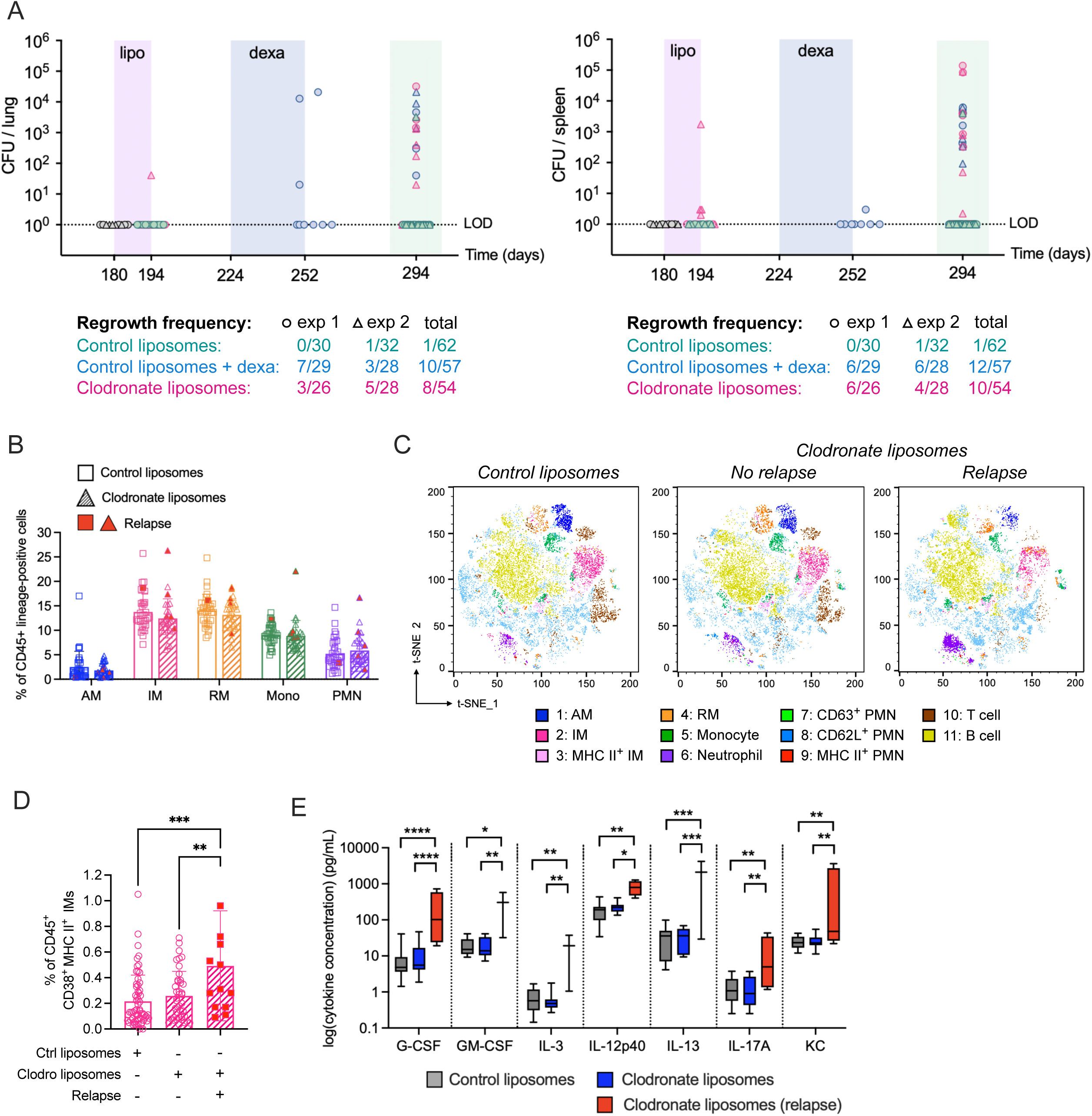
Depletion of interstitial macrophages during latency results in increased tuberculosis relapse in mice, accompanied by elevated pro-inflammatory interstitial macrophages and cytokines in the lung. (A) Bacterial burden prior and after IM depletion was assessed by CFU determinations in the lung and spleen of mice. Different symbols (circle, triangle) denote date from two independent experiments that were combined for analysis. Numbers below each graph indicate the frequency of Mtb regrowth in mice treated with control liposomes with or without dexa, or with clodronate liposomes. (B) Immunoprofiling performed by flow cytometry three months post-depletion to assess the frequency of each immune cell types in the lung of mice treated with control and clodronate liposomes, where the red symbols indicate mice that relapsed. (C) t-SNE plots of cell subtypes after clustering, colored according to cellular identity and separated by condition immediately following treatment, namely control liposomes, and mice treated with clodronate liposomes with or without relapse. (D) Frequency of pro-inflammatory Ims (CD38^+^ MHC II^+^) present in the lung of treated mice three-months post-depletion as assessed by flow cytometry. (E) Cytokine concentrations in lung homogenates from mice treated with control liposomes (gray), or from clodronate liposome-treated mice that relapsed (red) or did not relapse (blue). Data represents the mean ± SD of 4 to 32 biological replicates. * p < 0.05, ** p < 0.01, *** p < 0.001, **** p < 0.0001 based on Poisson test (for A), two-way ANOVA (for B) or one-way ANOVA (for D) statistical test with Tukey’s multiple comparisons test or an F-test (for E).

The Poisson model was used for evaluating differences in reactivation rates (**Figure 3A**). To estimate the reactivation rates for each treatment, data from mice at multiple timepoints was combined by summing the number of mice at each timepoint weighted by the number of days, i.e. “mouse-days”. Data was combined over two independent experiments. We used a Poisson test to quantify the statistical significance of differences among these rates of reactivation. Specifically, we applied an exact test of the ratio of Poisson rate parameters (Gu et al., 2008), as implemented in the *poisson.test()* function in R.

## 3. RESULTS

### The pulmonary myeloid cell compartment and cytokine milieu are altered in a paucibacillary mouse model of latent tuberculosis infection

Considering the enhanced ability of IMs to restrict Mtb during acute infection, we sought to study their importance in controlling persisting Mtb during latency in mice. We adapted our previously described paucibacillary mouse model of LTBI (H. Su et al., 2021) by adding a two-week treatment with clodronate-loaded or control liposomes during the latent phase to assess the importance of IMs in TB relapse (**Figure 1A**). As in our previous studies, Mtb BPL-DUC replicated during the first month of infection and was undetectable in lungs and spleens following a 16-week doxycycline treatment due to the depletion of BPL causing death of Mtb (**Figure 1B**).

We characterized the pulmonary immune cell landscape by flow cytometry throughout the course of the experiment and identified alveolar macrophages (AMs: Ly6G^-^ CD11b^-^ CD11c^+^ Siglec-F^+^), interstitial macrophages (IMs: Ly6G^-^ CD11b^+^ Siglec-F^-^ CX3CR1^+^), recruited monocytes and macrophages (RMs: Ly6G^-^ CD11b^+^ Ly6C^-^), monocytes (Monos: Ly6G^-^ CD11b^+^ Ly6C^+^) and neutrophils (PMNs: Ly6G^+^ CD11b^+^) from CD45^+^ cells, as described in our gating strategy (**Supp. Figure 1**). At the peak of infection on day 28, we found the largest proportions of cells in the lung were IMs and monocytes (**Figure 1C**), while RMs were the predominant immune cell type during the latent phase of infection on day 180 (**Figure 1D**). Analyzing the changes of cell population frequencies over time revealed that AMs and neutrophils remained at relatively low levels throughout the infection, while IMs and monocytes increased during the first 4 weeks of infection and decreased to baseline levels alongside the elimination of Mtb. In contrast, RMs were continuously recruited and the most abundant immune cell type during latency (**Figure 1E**). These changes in the myeloid cell compartment are also evident in t-SNE plots visualizing each marker used in our flow cytometry panel. At the peak of infection, we identified an increased number of cells expressing CD11b, MHC II, CD38 and Ly6C, consistent with the highest frequency of IMs and monocytes at this time point (**Supp. Figure 2A**). On the other hand, fewer cells expressed these markers during the paucibacillary phase of infection (**Supp. Figure 2B**).

To assess the activity of these macrophage and neutrophil populations, we measured the concentrations of various cytokines in the lungs of mice at day 1, day 28 and day 180 of the infection. The levels of several pro-inflammatory cytokines were increased at the peak of infection at day 28 compared to uninfected mice (**Figure 1F**). The elevated levels of IL-6, IL-10 and TNF were sustained during latency at day 180. In contrast, the levels of IL-1β, MIP-1ɑ, MIP-1β and RANTES decreased to baseline during the latent phase. IL-12p70 was the sole cytokine measured that increased in the lung of mice during latency compared to peak infection (**Figure 1F**). These data suggest that a baseline level of inflammation is necessary to sustain control of Mtb during latency, while simultaneously limiting further immune cell recruitment.

### Clodronate liposome treatment during latency leads to a depletion of interstitial and recruited macrophages, and increased recruitment of neutrophils to the lung

Having established the paucibacillary, latent phase of TB infection, we treated infected mice with control or clodronate liposomes for two weeks to deplete IMs (**Figure 1A**). We performed immunoprofiling of the myeloid cell compartment in the lung (**Figure 2A,C**), spleen (**Figure 2D-F**) and blood (**Figure 2G-I**) immediately post-depletion. As expected, in the lung we observed a significant decrease in the number of IMs and RMs in mice treated with clodronate liposomes compared to those treated with control liposomes (**Figure 2A**). In contrast, AMs were not significantly affected by the intravenous administration of clodronate liposomes. Surprisingly, we observed a two-fold increase in the number of neutrophils recruited to the lung upon IM depletion (**Figure 2A**). One week after depletion of IMs, all monocyte, macrophage and neutrophil populations had regained homeostasis, as indicated by the similar cell population frequencies between mice treated with control or clodronate liposomes (**Figure 2B**). The transitory neutrophils displayed no difference in their activation markers CD62L and MHC II or in the degranulation marker CD63 whether mice were treated with control or clodronate liposomes (**Figure 2C**). Since neutrophils were recruited to the lung following IM depletion, we assessed their abundance, activation and functionality in the spleen and blood as well. We observed by immunoprofiling that although the number of neutrophils in the spleen of infected mice was reduced during latency compared to uninfected mice, there was no difference whether IMs were depleted or not (**Figure 2D**). However, in mice depleted of IMs the frequency of CD62L^+^ neutrophils was increased (**Figure 2E**) and the frequency of MHC II^+^ neutrophils was decreased (**Figure 2F**) compared to control mice. These data are consistent with a shift towards a more immature or less activated splenic neutrophil population upon IM depletion. We found no difference in the proportion of neutrophils in uninfected mice or infected mice treated with either liposome treatment in the circulation as assessed by immunoprofiling blood (**Figure 2G**). To investigate the functionality of these neutrophils, we quantified reactive oxygen species (ROS) production which revealed no difference in the frequency of neutrophils that produce ROS (**Figure 2H**) or in the amount of ROS produced by neutrophils (**Figure 2I**), whether they were treated with control or clodronate liposomes. These data indicate that neutrophils isolated from IM-depleted mice were as functionally active as those from control mice. We simultaneously characterized dendritic cell populations in the lung and spleen of these mice immediately following IM depletion. We found no difference in the frequency of CD24^+^ CD11c^+^ pulmonary and splenic DCs, but CD11c^+^ DCs expressed more CD24 when IMs were absent (**Supp. Figure 3A, C**). Clodronate liposome-treated mice also had an increased frequency of CD103^+^ CD11c^+^ DCs in the lung (**Supp. Figure 3B**), but not in the spleen (**Supp. Figure 3D**). These data suggest a potential shift toward cross-presenting, migratory cDC1 in the lung after IM depletion, which could aid in tighter bacterial control or limiting excessive inflammation. Additional studies would be required to validate these findings.

We subsequently performed high dimensionality reduction and clustering analysis to visualize the remodeled immune landscape during latency following IM depletion. We obtained 12 main cell clusters based on our immune cell flow cytometry panel, as demonstrated by the t-SNE plots (**Figure 2J**). In infected mice that were treated with control liposomes, IM (2) and RM (4) cell clusters were enriched compared to uninfected mice, and these populations were almost entirely absent in mice treated with clodronate liposomes. There were also more cells in the neutrophil cluster (6) upon clodronate treatment compared to control (**Figure 2J**). This high dimensionality reduction analysis highlights the immune landscape reprogramming observed in mice depleted of IMs, with a reduction of IMs and RMs, and an increase in neutrophils in the lung immediately following IM depletion. Additionally, the t-SNE plots for each flow cytometry marker reveal the increased number of cells expressing Ly6G compared to pre-depletion, consistent with the increased recruitment of neutrophils to the lung (**Supp. Figure 2C**). We further assessed the activity of these cells by determining the levels of various cytokines in the lungs of these mice. Of all the cytokines measured, we found that only the abundance KC and MIP-1ɑ was elevated in the lung of mice treated with clodronate liposomes (**Figure 2K**), while all other cytokines analyzed had comparable levels in mice treated with control or clodronate liposomes (**Supp. Figure 4**).

### Depletion of interstitial macrophages during latency leads to increased tuberculosis relapse accompanied by elevated pro-inflammatory interstitial macrophages and cytokines in the lung

Throughout the course of the chronic infection, we assessed bacterial burden in the lung and spleen of mice. As expected, we observed bacterial regrowth in mice immunosuppressed with dexamethasone in the lung and spleen corresponding to a 28% relapse frequency compared to 2% relapse of mice treated with control liposomes (**Figure 3A**, **Table 1**). In mice that were depleted of IMs with clodronate liposomes, Mtb reactivated in the lung and spleen, corresponding to a frequency of 26% TB relapse (**Table 1**). These data demonstrate that transient depletion of IMs during the paucibacillary phase phenocopies the effect of dexamethasone-induced immunosuppression, leading to enhanced TB relapse.

**Table 1.** Analysis of Mtb regrowth and relapse frequencies. Data from two independent experiments including number of mice, frequency of regrowth in the lung and in the spleen as assessed by colony forming units (CFUs), number of mice that relapsed, frequency of relapse and statistics. The Poisson model was used to evaluate statistical differences in reactivation rates.

|  | **Control liposomes** | **Dexamethasone-induced** | **Clodronate liposomes** |
| --- | --- | --- | --- |
| Experiments | 2 | 2 | 2 |
| Mice | 62 | 57 | 54 |
| Mice with regrowth in the lung (CFU ≥ 1) | 1 | 10 | 8 |
| Mice with regrowth in the spleen (CFU ≥ 1) | 1 | 12 | 10 |
| Relapsed, CFU in lung or spleen | 1 | 16 | 14 |
| Relapse frequency (%) | 1.6 | 28.1 | 25.9 |
| p value, compared to control | n/a | 2.13E-07 | 2.73E-04 |
| p value, compared to dexamethasone | 2.13E-07 | n/a | > 0.05 |

To understand the dynamic nature of the myeloid cell compartment in our model, we characterized the immune cells at endpoint, 10-months post-infection, by immunoprofiling the pulmonary myeloid cells by flow cytometry. We observed that three months after liposome depletion, all myeloid populations assessed, AMs, IMs, RMs, monocytes and neutrophils, had returned to baseline levels pre-depletion (**Figure 3B**). Comparing t-SNE plots of mice treated with control liposomes with those of clodronate liposomes-treated mice that relapsed or did not relapse, we did not observe major changes in the cell clusters (**Figure 3C**), emphasizing the return to homeostasis of the analyzed immune cell populations. Segregating MHC II^+^ CD38^+^ pro-inflammatory IMs by TB disease state revealed a significantly higher proportion of these in mice that relapsed compared to mice that did not relapse, or to control mice (**Figure 3D**). Furthermore, measuring the concentration of various cytokines expressed in the lungs of these mice three months post-depletion revealed that G-CSF, GM-CSF, IL-3, IL-12p40, IL-13, IL-17A and KC were elevated in lungs of mice that relapsed compared to mice that did not relapse and to control mice (**Figure 3E**), indicating that Mtb reactivation is associated with a reprogramming of the pulmonary cytokine milieu. t-SNE plots for each immune cell marker at this time point showed little difference in cell numbers compared to the previous timepoint post-depletion (**Supp. Figure 2C,D**).

### Spleen pathology in mice treated with clodronate liposomes does not correlate with Mtb burden or TB relapse

Unexpectedly, at endpoint, three months post-depletion of IMs, we observed that most spleens from clodronate liposome-treated mice appeared fibrotic or necrotic with white lesions partially or completely penetrating the spleen (**Figure 4A**). In fact, 68% of clodronate liposome-treated mice displayed such pathology compared to 0% in control mice (**Figure 4B**). In addition, these spleens were very rigid and difficult to homogenize, and most of the cells collected from these spleens were dead, as assessed by flow cytometry (**Figure 4C**). As a result, immunoprofiling of splenocytes from these mice was not possible at this time point. Importantly, we found no correlation between Mtb regrowth in the spleen and the extent of necrosis (**Figure 4D**), or between the mice that had TB relapse and the extent of necrosis of the spleen (**Figure 4E**). Thus, while most mice treated with clodronate liposomes suffered from splenic necrosis, this pathology does not correlate with Mtb regrowth in the spleen or with TB relapse.

**Figure 4.**
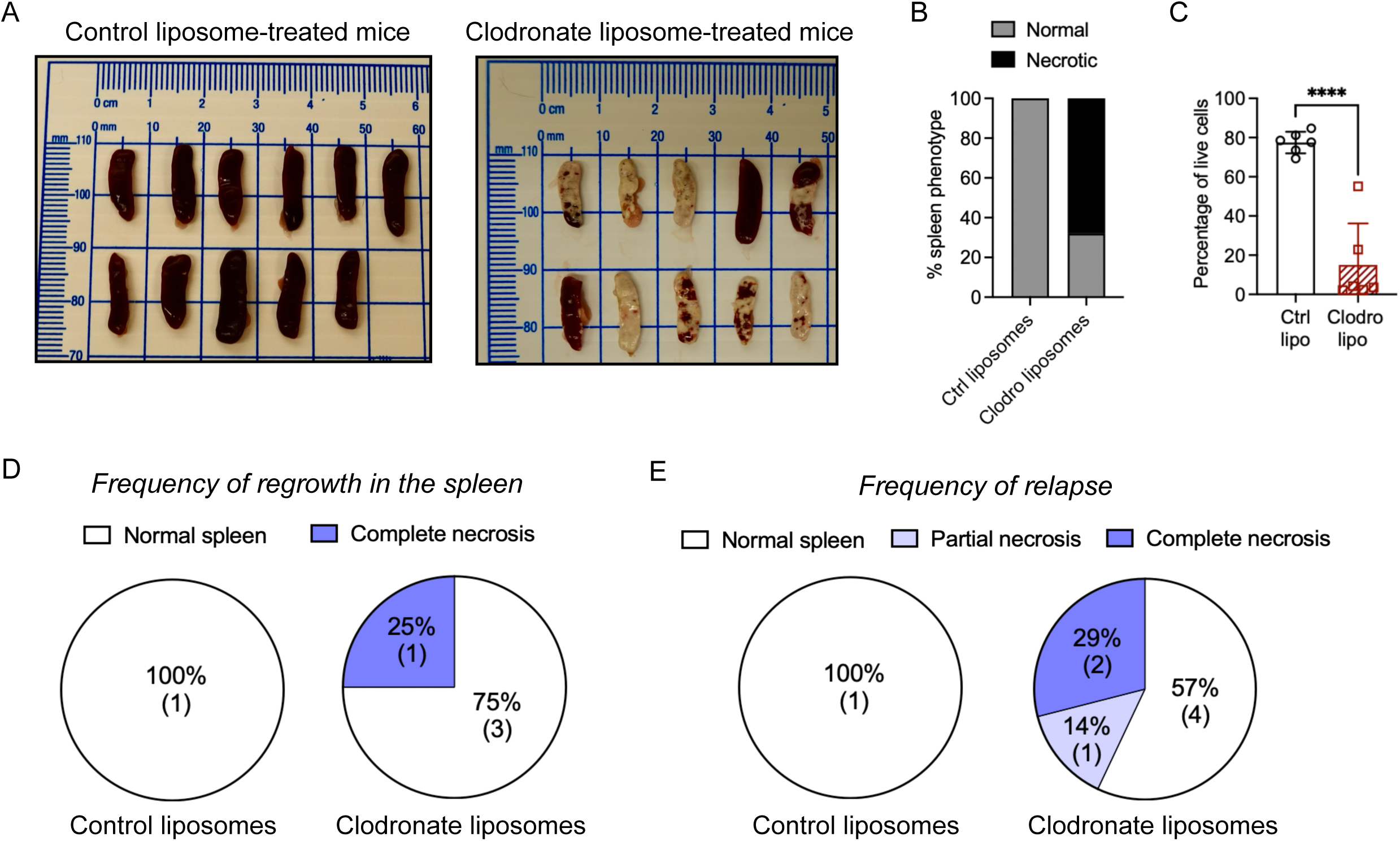
Spleen pathology in mice treated with clodronate liposomes does not correlate with Mtb bacterial burden or TB relapse. (A) Gross pathology of spleens harvested from control liposome-or clodronate liposome-treated mice three months post-depletion. (B) Percentage of spleens that appear necrotic compared to normal in the two treatment groups. (C) Flow cytometry was performed with splenocytes isolated from control liposome-treated (Ctrl lipo) and clodronate liposome-treated (Clodro lipo) mice at endpoint and the percentage of live cells was assessed. (D-E) Frequency of Mtb regrowth in the spleen (D) and frequency of relapse (E) of control liposome- and clodronate liposomes-treated mice segregated by degree of necrosis of the spleen. Number in bracket indicates the number of mice.

## 4. DISCUSSION

To uncover the host mechanisms that allow for control of Mtb and to elucidate the role of the innate immune system in preventing transition from latency to active TB disease, we used our previously established paucibacillary mouse model of latency (H. Su et al., 2021). We discovered that IMs are essential in restricting Mtb and preventing Mtb reactivation and disease. These results corroborate our prior assumption that BPL depletion would synergize with host innate immunity to control Mtb (H. Su et al., 2021; Tiwari et al., 2018). Although this mouse model does not perfectly replicate latency in humans, it can be used as a tool to explore host-derived factors and enrich our understanding of human TB based on the similar paucibacillary infection states during latency and relapse between these hosts.

At the peak of infection with Mtb (day 28), IMs, RMs and monocytes were the predominant components of the pulmonary myeloid cell landscape, in contrast to AMs and neutrophils which remained at relatively low levels at this stage of the infection (**Figure 1C,E**). During the paucibacillary phase, the RM population expanded as the requirement to support and replenish pre-existing IMs arose (**Figure 1D,E**). Throughout the course of latent infection prior to the depletion of IMs, we observed an associated remodeling of the cytokine environment (**Figure 1F**), presumably in response to the Mtb load present at these various timepoints and to fine-tune the immune response to this inflammatory insult. Various cytokines, notably IL-6, IL-10 and TNF, were increased at the peak of infection and their levels maintained throughout latency, where IL-6 and TNF had the largest increase in concentration compared to basal levels at day 1 post-infection (**Figure 1F**). Immediately following IM depletion, most cytokines were comparable between mice with depleted IMs and controls. However, we observed approximately 5-fold elevated levels of KC and MIP-1ɑ in the lung of IM-depleted mice compared to control mice (**Figure 2K**). Both KC and MIP-1ɑ are potent neutrophil chemoattractants (De Filippo et al., 2008; Hilda et al., 2014; Johnson et al., 2011; Ritzman et al., 2010; Tumpey et al., 2002), while MIP-1ɑ also promotes the recruitment of monocytes, T cells and dendritic cells (DCs) via secretion from activated macrophages (Bhavsar et al., 2015; Madsen et al., 2003). MIP-1ɑ not only boosts macrophage phagocytosis and microbial killing but also aids in coordinating lesion formation to contain Mtb (Saukkonen et al., 2002; Silva Miranda et al., 2012). This is likely a compensatory mechanism by other myeloid cells in the lung to counterbalance the absence of IMs at that time. As we had hypothesized, depletion of IMs during the latent phase of this model led to a heightened bacterial burden in the lung and spleen of mice three months post-depletion, corresponding to an increased frequency of TB relapse in these mice (**Figure 3A**, **Table 1**). At this time, we observed an enrichment of pro-inflammatory MHC II^+^ CD38^+^ IMs in mice that relapsed (**Figure 3D**). Both MHC II and CD38 have been shown previously to be biomarkers on IFN-γ^+^ CD4^+^ T cells that discriminates between active TB and LTBI in two cohorts (Balcells et al., 2019; Dutta et al., 2014; Flynn et al., 2011a; McCaffrey et al., 2022; Silva Miranda et al., 2012). It wil be interesting to investigate in future studies whether this could also be the case for IMs.

At the final time point, we observed an increase of G-CSF, GM-CSF, IL-3, IL-12p40, IL-13, IL-17A and KC secreted in the lung of clodronate liposome-treated mice that relapsed (**Figure 3E**) compared to mice that did not relapse and to control mice. The elevated levels of these cytokines typically reflect an enhanced pro-inflammatory response that largely promotes macrophage (GM-CSF, IL-12p40) and neutrophil (G-CSF, GM-CSF, IL-17A, KC) recruitment and activation (Bajrami et al., 2016; Castellani et al., 2019; De Filippo et al., 2008; Griffin et al., 2012; Ha et al., 1999; Kumar et al., 2022; K. M. C. Lee et al., 2020; Ritzman et al., 2010). Elevated IL-12p40 and IL-17A levels during TB relapse without an associated increase in IL-12p70 indicate a shift from Th1 immunity toward Th17 responses (Khader & Cooper, 2008; Lyakh et al., 2008). The associated changes in the cytokine profiles may contribute to, or correlate with TB relapse.

The increased frequency of neutrophils in the lungs immediately following IM depletion was unexpected (**Figure 2A**). A recent study revealed that the use of clodronate liposomes in a model of serum transfer arthritis led to stunning of neutrophils, a transient state of functional unresponsiveness or suppression following strong activation, such as in response to severe infection or inflammation (Culemann et al., 2023; Leliefeld et al., 2016). The authors showed that administering clodronate liposomes to mice led to a block of neutrophil phagocytic capacity, cytokine expression, ROS production and altered their migratory and swarming behavior (Culemann et al., 2023). Because we observed an increased recruitment of neutrophils to the lung upon depletion of IMs with clodronate liposomes in our study, we assessed the phenotype and functionality of these neutrophils. We demonstrated that the neutrophils recruited to the lung of clodronate liposome-treated mice in our model of Mtb latent infection were functionally similar to those in control mice, as assessed by their comparable expression of CD62L, MHC II and CD63 in the lung (**Figure 2C**). In the spleen, there was a higher frequency of CD62L^+^ neutrophils (**Figure 2E**), suggesting either a reduced activation state or a transitional phenotype (Drescher et al., 2020; Leliefeld et al., 2016; Malengier-Devlies et al., 2021; Paudel et al., 2022), whereas the observed decrease in MHC II^+^ splenic neutrophils suggests reduced maturation (Borkute et al., 2021; Forrer et al., 2024; Vono et al., 2017). IM depletion may lead to systemic immune dysregulation, triggering emergency granulopoiesis or systemic inflammation (Bain & MacDonald, 2022; Bedoret et al., 2009; Hou et al., 2021; Tamoutounour et al., 2013; Zaynagetdinov et al., 2013). It is possible that IMs indirectly regulate neutrophil maturation, so their depletion may lead to a loss of immunoregulatory cues and to the observed phenotypes (Dang et al., 2023; Maier-Begandt et al., 2024; Y. Su et al., 2020). We further characterized the neutrophil function by assessing the production of ROS by circulating neutrophils and found no difference amongst mice treated with control or clodronate liposomes (**Figure 2G-I**). These data together suggest that these neutrophils are most likely not functionally impaired or “stunned” upon depletion of IMs at a systemic level but might be more immature in the spleen. As mentioned, the elevated expression of the neutrophil chemoattractants KC and MIP-1ɑ in the lungs of mice treated with clodronate liposomes (**Figure 2K**) is consistent with the increased neutrophil recruitment observed at this inflammatory site (De Filippo et al., 2008; Hilda et al., 2014; Ritzman et al., 2010).

Interestingly, CD62L^+^ MHC II^lo^ neutrophils have been shown to mediate tissue damage in different mouse models (Drescher et al., 2020; Ling & Xu, 2024), which we observed at endpoint. The spleens from mice depleted of IMs three months prior had undergone partial or complete cell death in the form of fibrosis or necrosis (**Figure 4A**), indicating impaired immune regulation or unresolved inflammation. At this time, we observed an increased production of IL-13 in the lung of mice that relapsed (**Figure 3E**). IL-13 is a Th2 cytokine that typically dampens classical macrophage killing pathways, promotes fibrosis and destabilizes the granuloma structure, enabling dissemination and relapse (Cronan, 2022; Flynn et al., 2011b; C. G. Lee et al., 2001; Walter et al., 2022). The tissue remodeling environment promoted by IL-13 can undermine bacillary control (Asquith et al., 2011; Heitmann et al., 2014). On the other hand, the upregulated secretion of IL-17A in the lung of mice with TB relapse, particularly alongside elevated IL-12p40 but not IL-12p70 levels (**Figure 3E**), may indicate a shift towards a more IL-17-driven inflammatory milieu resulting from the failure of IMs to sterilize the bacilli during latency when they are depleted, ultimately triggering tissue damage and Mtb reactivation. Together, these data point to a model where abruptly depleting IMs during latency leads to the loss of Mtb’s key intracellular niche. Damage-associated molecular patterns (DAMPs) and bacterial components trigger the production of KC and MIP-1ɑ, perhaps activating emergency granulopoiesis in the spleen and ultimately leading to TB relapse following a shift in the cytokine milieu that favors an IL-17A-driven response.

## ACKNOWLEDGEMENTS

We thank Sae Woong Park, Kaj Kreutzfeldt and Xiuju Jiang for technical assistance, and Kathryn Dupnik, Rachel Kinsella, Christina Stallings and Chen Yu for advice and helpful discussions. This work was supported by the Tri-Institutional TB Research Unit (National Institutes of Health grant U19 AI111143). V. Vinette was a recipient of a Potts Memorial Foundation Grant and an American Association of University Women (AAUW) International Fellowship. Author contributions: V. Vinette, M. Gengenbacher and S. Ehrt conceived the study and analyzed the data; V. Vinette, A. Castro, H. Kim, C. Trujillo and M. Xie performed the experiments; V. Vinette and T.R. Ioerger performed statistical analyses; V. Vinette and S. Ehrt wrote the manuscript with input from the other authors.

