## Supplementary material for "Interstitial macrophages prevent tuberculosis relapse by restricting *Mycobacterium tuberculosis* immune evasion"

**Supplementary Figure 1 – Gating strategy to identify myeloid cell populations.** Gating strategy to identify neutrophils (PMNs, purple), alveolar macrophages (AMs, blue), recruited monocytes/macrophages (RMs, orange), monocytes (Monos, green) and interstitial macrophages (IMs, pink) by flow cytometry.

**Supplementary Figure 2 – Temporal dynamics of immune cell marker expression.** High dimensionality reduction and clustering analyses were performed from datasets acquired by flow cytometry throughout the course of the experiment. FlowSOM clustering was performed on t-distributed stochastic neighbor embedding (t-SNE) plots to visualize the markers used to characterize various immune cell type at (A) peak of infection, (B) during the latency phase, (C) after depletion of interstitial macrophages, and (D) at experimental endpoint, three months post-depletion.

**Supplementary Figure 3 – Immunophenotype of dendritic cells in the lung and spleen upon interstitial macrophage depletion during latent tuberculosis infection.** Lung and spleen were harvested from mice treated with control or clodronate liposomes immediately following depletion of interstitial macrophages. Immunoprofiling was performed by flow cytometry to assess pulmonary $CD11c^+$ DCs and their frequency and gMFI of CD24 (A) and CD103 (B), as well as the frequency of splenic $CD11c^+$ DCs for CD24 (C) and CD103 (D). Data represents mean ± SD of 5 biological replicates. * $p < 0.05$ based on t-test statistical test.

**Supplementary Figure 4 – Pulmonary cytokine levels after treatment with liposomes.** Quantification of cytokine concentrations in the lung of mice treated with control or clodronate liposomes immediately following liposome treatment. Data represents the mean ± SD of three to five biological replicates. Statistical analysis performed by two-way ANOVA statistical test with Tukey’s multiple comparisons test.

Supp. Figure 1 - Gating strategy to identify myeloid cell populations

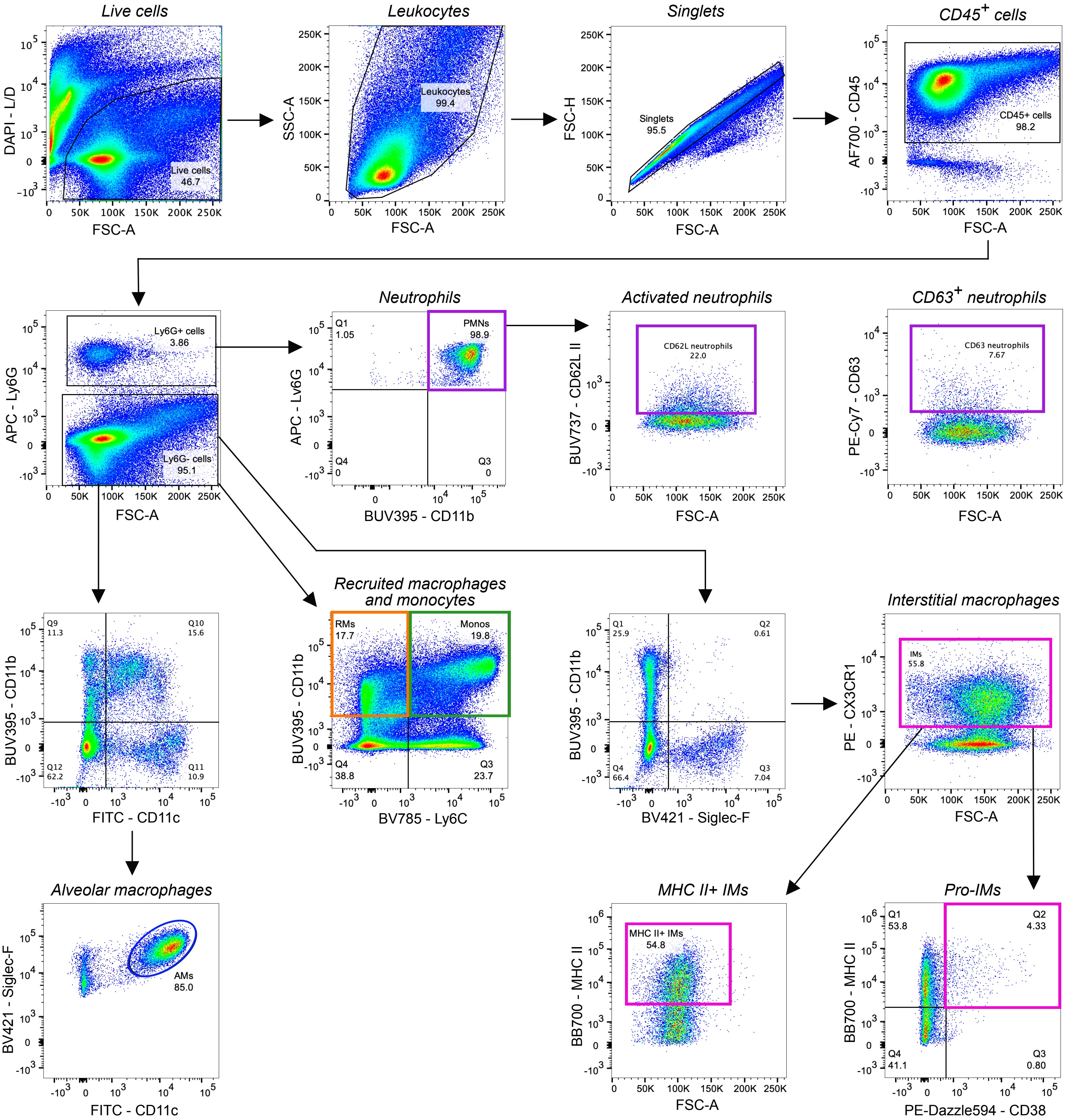

Supp. Figure 2 - Temporal dynamics of immune cell marker expression

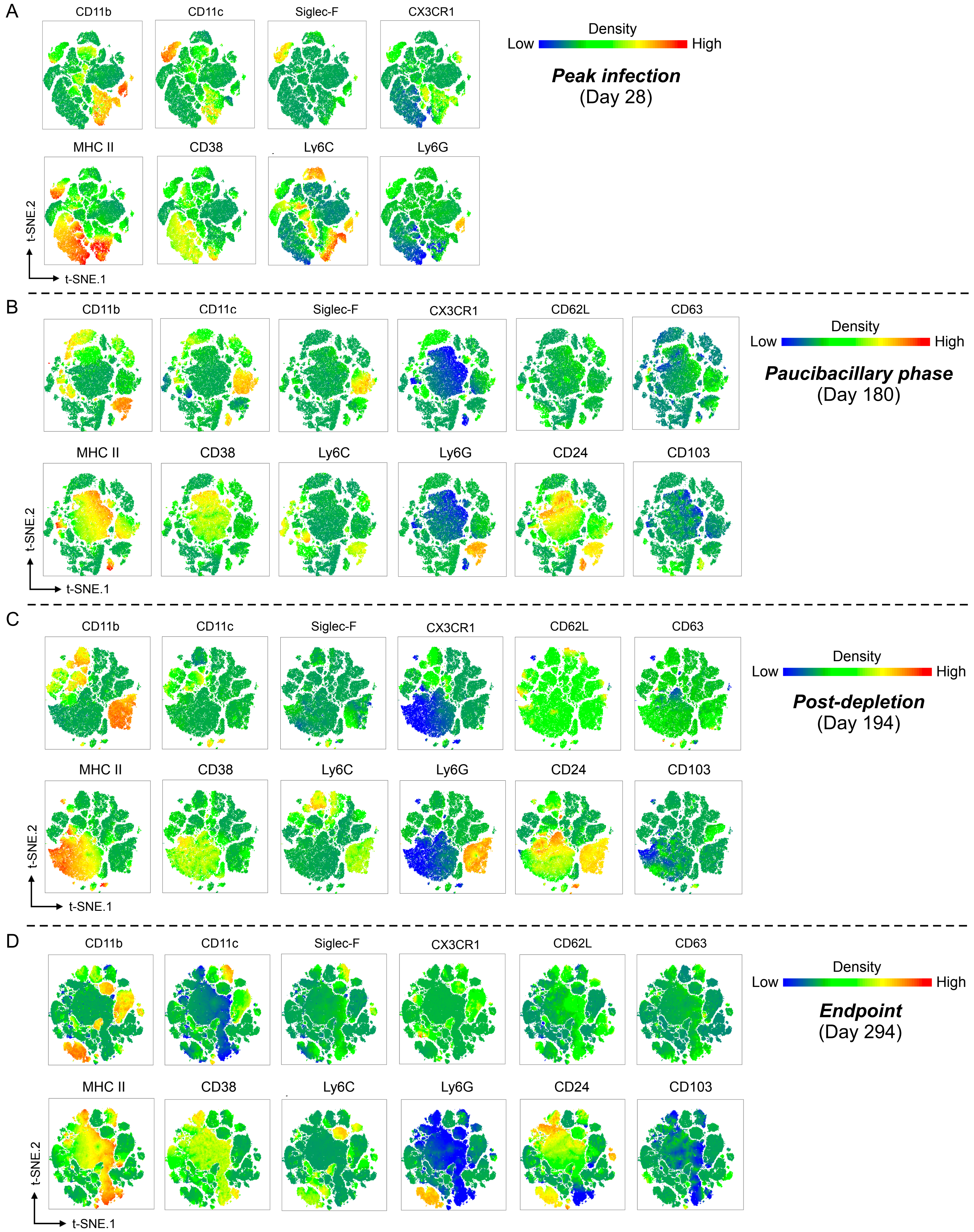

Supp. Figure 3 - Immunophenotype of dendritic cells in the lung and spleen upon interstitial macrophage depletion during latent tuberculosis infection

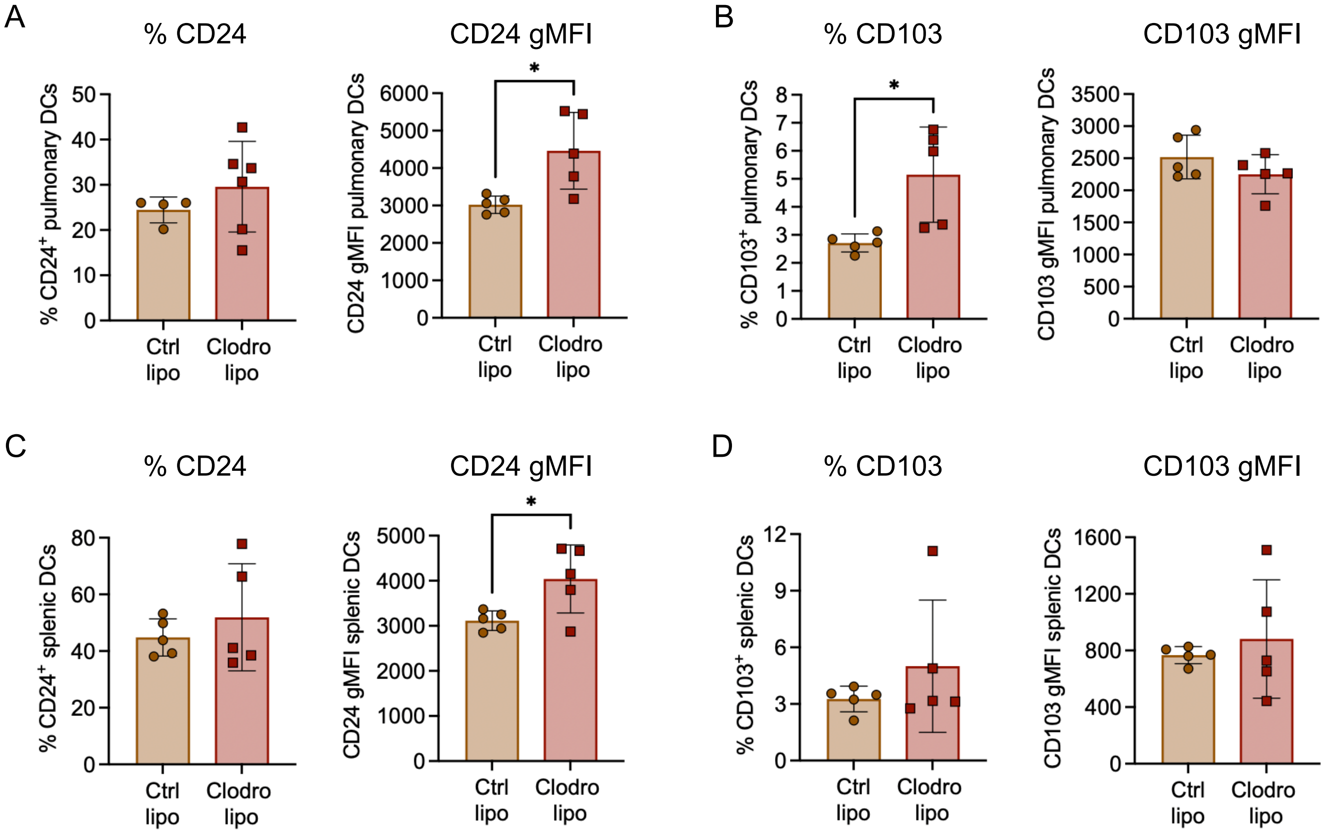

Supp. Figure 4 - Pulmonary cytokine levels after treatment with liposomes

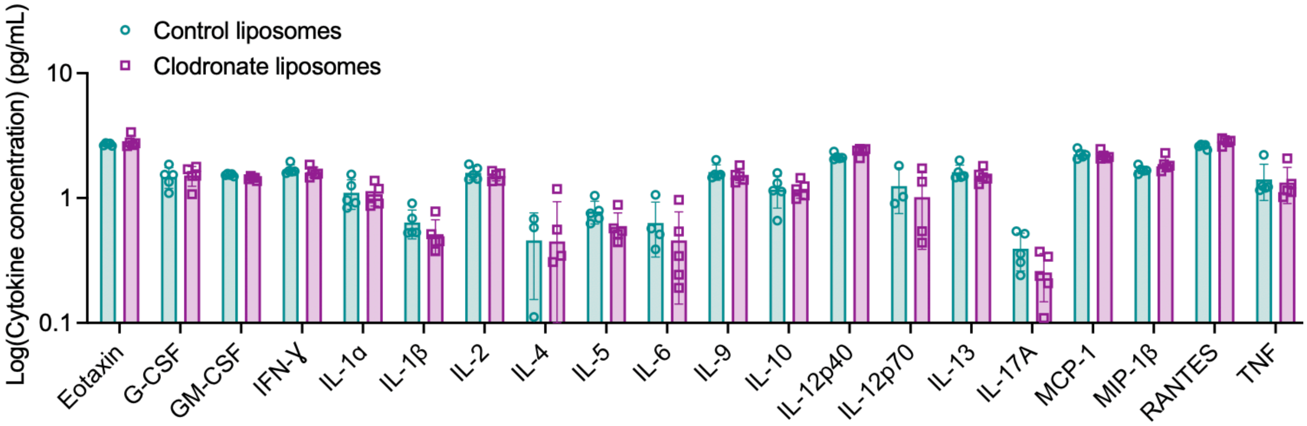
